# Efficient evidence-based genome annotation with EviAnn

**DOI:** 10.1101/2025.05.07.652745

**Authors:** Aleksey V. Zimin, Daniela Puiu, Mihaela Pertea, James A. Yorke, Steven L. Salzberg

## Abstract

For many years, machine learning-based *ab initio* gene finding approaches have been central components of eukaryotic genome annotation pipelines, and they remain so today. The reliance on these approaches was originally sustained by the high cost and low availability of gene expression data, a primary source of evidence for gene annotation along with protein homology. However, innovations in modern sequencing technologies have revolutionized the acquisition of gene expression data, allowing scientists to rely more heavily on this class of evidence. In addition, proteins found in a multitude of well-annotated genomes represent another invaluable resource for gene annotation. Existing annotation packages often underutilize these data sources, which prompted us to develop EviAnn (<u>Evi</u>dence-based <u>Ann</u>otator), a novel evidence-based eukaryotic gene annotation system. EviAnn takes a strongly data-driven approach, building the exon-intron structure of genes from transcript alignments or protein-sequence homology rather than from purely *ab initio* gene finding techniques. We show that when provided with the same input data, EviAnn consistently outperforms current state-of-the-art packages including BRAKER3, MAKER2, and FINDER, while utilizing considerably less computer time. Annotation of a mammalian genome can be completed in less than an hour on a single multi-core server. EviAnn is freely available under an open-source license from https://github.com/alekseyzimin/EviAnn_release and from Bioconda as “eviann”.

## Introduction

Two of the most reliable sources of evidence for gene annotation are gene expression data and protein homology. Gene expression data can be captured with high-throughput RNA sequencing (RNA-seq) technology, which sequences all the genes being expressed in a tissue sample and can easily generate tens of millions of reads per sample. These short reads can be aligned to a genome and then assembled to create an accurate picture of the exon-intron structures of all the genes and splicing variants present in the original tissue. RNA-seq technology has been augmented in recent years with the introduction of long-read sequencing and even direct sequencing of RNA, which can capture full-length transcripts and thereby provide even more accurate exon-intron structures. However, RNA-seq technology is far from perfect: most RNA-seq protocols do not recover complete transcripts, meaning that the sequence fragments need to be assembled, which in turn may introduce errors. In addition, an RNA-seq experiment only measures the genes expressed in one tissue (or one cell, in some cases), and thus RNA from multiple tissues and conditions is needed to capture all genes and transcripts for a comprehensive annotation. Along with these advances in transcriptome sequencing, the scientific community has now assembled and annotated genomes for thousands of different species. These genomes provide an additional valuable source of annotation evidence, particularly in the form of protein sequences that are well conserved even among distant species.

Most of the current genome annotation systems, including MAKER2 (Cantarel et al., 2008; Holt and Yandell, 2011), BRAKER3 (Gabriel et al., 2024), GeneMark-ETP (Bruna et al., 2024), and FINDER (Banerjee et al., 2021), do not use transcript sequencing evidence directly to produce gene annotation. Instead, they use transcript and protein sequence evidence to train *ab initio* gene finders such as GeneMark, SNAP (Korf, 2004), or AUGUSTUS (Stanke et al., 2006; Hoff and Stanke, 2019). The annotation systems then use the predictions made by these computational gene finders as the basis for their gene models. This approach was originally developed prior to the invention of next-generation RNA sequencing, at a time when expression data was expensive, few annotated genomes were available, and the number of known proteins was relatively small compared to today. The use of an *ab initio* gene finder could, in principle, compensate for gaps in RNA-sequencing data coverage by using machine-learning models to find genes at locations that are not expressed.

However, despite many years of algorithm development, *ab initio* gene finding is still highly error-prone for eukaryotic genomes, in which most of the sequence does not encode proteins. In particular, *ab initio* gene finders frequently generate very large numbers of false positive predictions, including erroneous exon-intron combinations and entirely erroneous gene loci. FINDER or BRAKER3 utilize protein homology from closely related species to filter and improve upon predictions made by their *ab initio* gene finders, but homology information cannot always fix all the errors. A further drawback is that *ab initio* gene finders typically only look for protein-coding sequences, which means they simply do not annotate untranslated regions (UTRs) of transcripts or long non-coding RNAs, the latter of which are a vital part of genome annotation today (Schuster and Hsieh, 2019; Cenik et al., 2020; Chatterjee et al., 2001).

In this paper, we describe a novel approach to genome annotation that relies primarily on direct evidence of expression or strong protein sequence similarity, avoiding the use of *ab initio* gene finding techniques. This approach is based on accurate processing and cross-referencing of different types of evidence, resulting in a faster and more transparent annotation process, where one can trace the origin of every annotated transcript or coding sequence (CDS) back to the input data. We have implemented this approach in an open-source software package called EviAnn (<u>Evi</u>dence-based <u>Ann</u>otator). EviAnn produces genome annotation by combining transcript assemblies created from RNA-seq data, transcripts from closely related species (if available), and protein sequences from at least one other related species. Because protein sequences are well conserved across the tree of life, if proteins from a closely related species are not available, EviAnn can instead use proteins from curated protein databases such as UniProt (Bairoch et al., 2005).

Below we compare EviAnn version 2.0.5 to current state-of-the-art annotation pipelines including MAKER2, BRAKER3, and FINDER on six plant and animal species: *Arabidopsis thaliana*, *Drosophila melanogaster*, *Populus trichocarpa* (poplar tree), *Danio rerio* (zebrafish), *Gallus gallus* (chicken), and *Mus musculus* (mouse). Our results demonstrate that, when running on the same data, EviAnn consistently produces gene annotations with superior accuracy and completeness, outperforming all other annotation pipelines. EviAnn also annotates many 5’ and 3’ UTR regions as well as long non-coding RNA genes that are missed by other methods. Finally, EviAnn runs very fast, more than ten times faster than all other pipelines.

## Results

The annotations for *A. thaliana*, *D. melanogaster*, and *M. musculus* have been extensively manually curated, and RefSeq versions of these annotations contain both a high-quality curated set of genes and transcripts and lower-confidence transcripts annotated by the NCBI Eukaryotic Genome Annotation Pipeline (https://www.ncbi.nlm.nih.gov/refseq/annotation_euk/process/). For our evaluations, we used the RefSeq identifiers to identify and exclude these lower-confidence transcripts (which start with the prefix XR for non-coding and XM for protein-coding transcripts) for these three species. The annotations for the remaining three species, *G. gallus*, *D. rerio,* and *P. trichocarpa*, were mostly or entirely produced by the NCBI pipeline. Thus, for these species, the performance metrics reported here are not necessarily reflective of the true accuracy of the pipelines; instead, they simply show how consistent the automated annotation pipelines are with the annotation produced by NCBI. EviAnn annotates both protein-coding genes and long non-coding RNAs (lncRNAs), and therefore we used reference annotations of both types of genes in this evaluation, as opposed to using only protein-coding genes, as has been done in some previous comparisons (Gabriel et al., 2024; Bruna et al., 2024).

We compared all annotation pipelines using a variety of metrics. First, as shown in Figure 1, we considered how accurately each method identified all genes in a genome, by which we mean the exon-intron boundaries as well as the transcription initiation and termination sites. Many genes have multiple splice variants (or *isoforms*), but for this evaluation, we only required a program to correctly annotate at least one isoform of each gene. For protein-coding genes, we required the pipeline to get the CDS boundaries correct, ignoring differences in noncoding exons. For noncoding RNA genes in the reference annotation, we required the pipeline to identify all introns correctly (from at least one isoform) to get credit for that gene. We allowed for some differences in the precise 5’ and 3’ boundaries of the transcripts (see Methods). As shown in the figure, EviAnn consistently demonstrated the highest accuracy on all six species tested, with particularly high precision (fewer false positives) on the chicken genome, *G. gallus*. Detailed values of sensitivity, precision, and F1 scores are given in Supplementary Tables S5–S7. These results show that even with a limited amount of RNA-seq data, EviAnn can produce the correct gene structure for ∼70% of high-confidence genes. Notably in this experiment, the output of StringTie2 alone, without further processing, was superior to both MAKER2 and FINDER.

**Figure 1.**
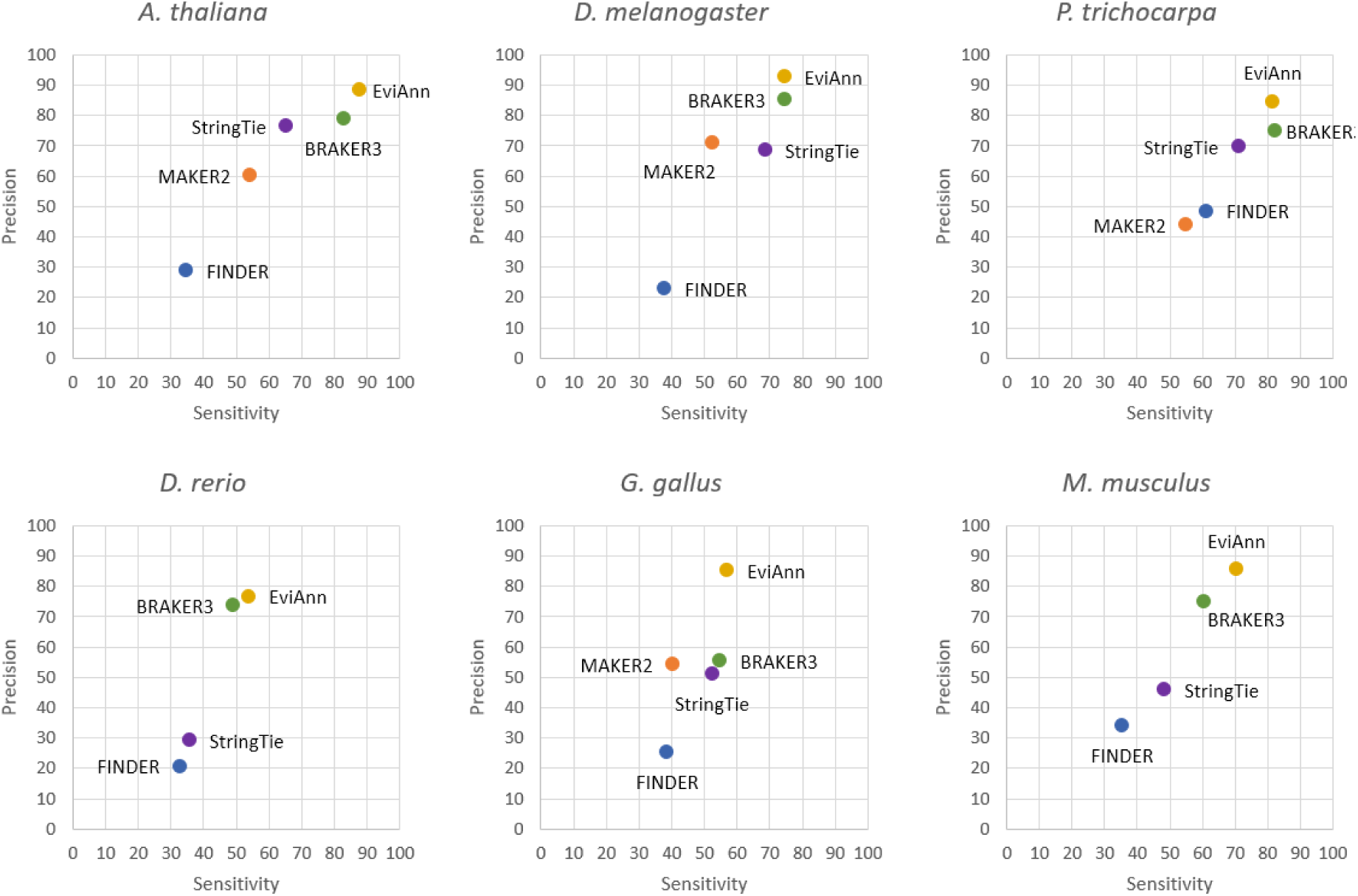
Accuracy of gene-level annotations by EviAnn, MAKER2, BRAKER3, and FINDER in six species. A gene locus was counted as correct if at least one transcript or CDS at that locus matched a reference transcript at the same locus, where a match required exact matches of all intron boundaries. Transcripts assembled with StringTie2 were used as input evidence for MAKER2, BRAKER3 and EviAnn. Data points representing the StringTie-assembled transcripts were added to illustrate how well those transcripts without further processing matched the references.

Our second evaluation focused on protein-coding sequences (CDSs). One reason for considering these separately is that some annotation pipelines, such as BRAKER3, only annotate the coding sequences in protein-coding genes, and do not attempt to find UTRs or noncoding RNA genes. For this comparison, we collected all unique coding sequences predicted by the annotation pipelines and matched them against all unique coding sequences present in the reference. The uniqueness criterion was important because pipelines such as MAKER2, FINDER, and EviAnn sometimes annotate multiple transcripts at a single locus that have the same coding sequence and that differ only in their noncoding regions.

Reference annotations can do the same; e.g., the RefSeq annotation for *D. melanogaster* has 27,284 protein-coding transcripts, but only 21,654 unique CDSs. In this evaluation, we counted a coding sequence as a match if the start codon, stop codon, and all intron boundaries within the coding regions matched the reference annotation exactly. Pipelines were not required to find noncoding exons or UTRs.

Results are shown in Figure 2. In this comparison, EviAnn again had the best overall performance, although BRAKER3’s statistics were nearly as good, with BRAKER3 getting slightly better precision and EviAnn showing slightly better sensitivity on average. As compared to the results at the gene level, overall sensitivity and precision were lower for the top-performing programs, likely because this second evaluation required programs to predict multiple protein-coding sequences at some loci.

**Figure 2.**
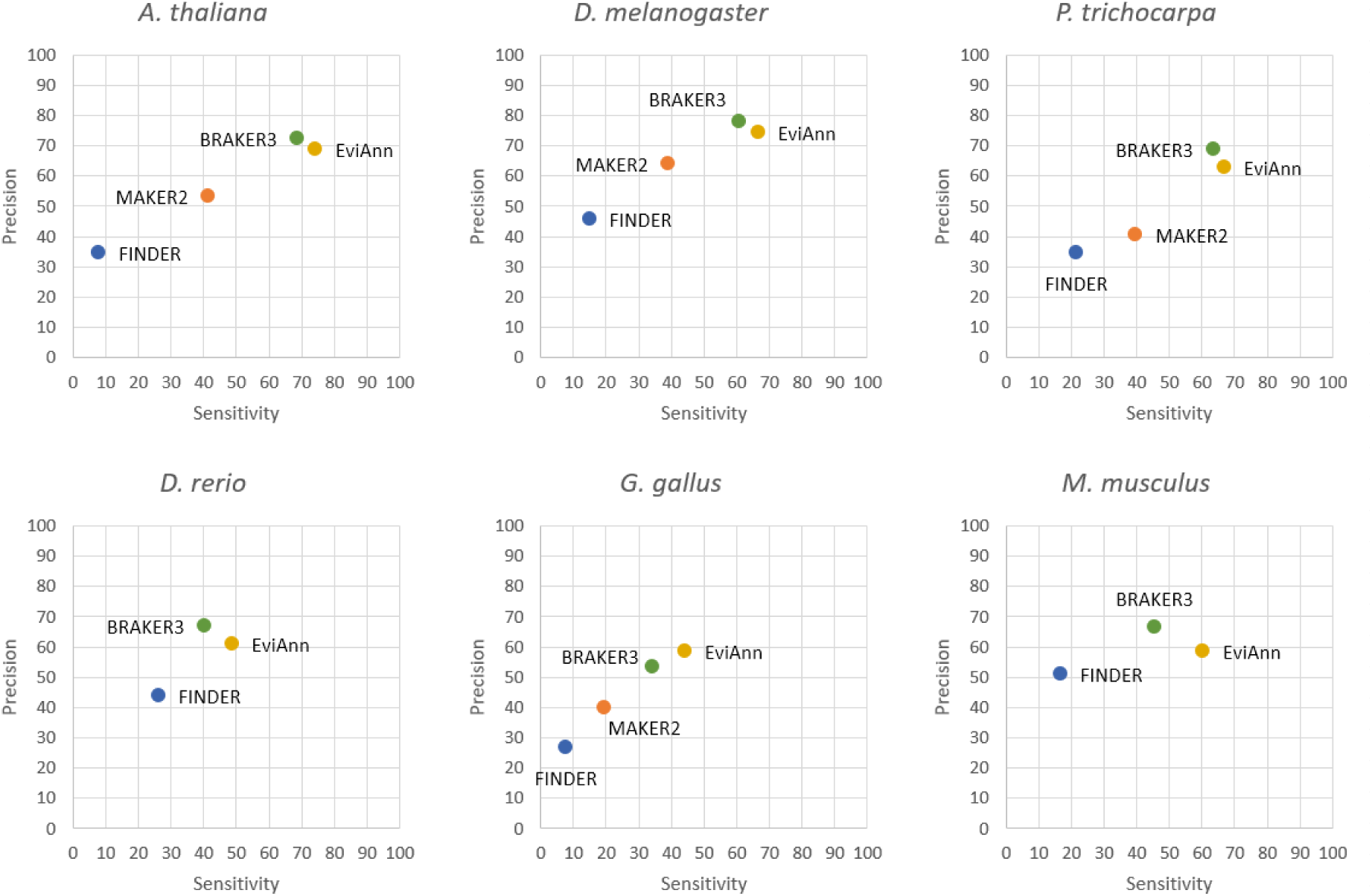
Accuracy of annotations of protein-coding sequences (CDSs) by EviAnn, MAKER2, BRAKER3, and FINDER in six species. A protein-coding annotation was considered correct if all coordinates of the protein-coding regions (CDSs) precisely matched the reference annotation. Noncoding exons were not considered in this evaluation.

Thirdly, we compared the pipelines on transcript level annotations, where we evaluated the number of reference transcripts that were matched by each pipeline (Figure 3). Here we counted all transcripts at every gene locus; i.e., if the reference annotation contained multiple splice variants for a given gene, then we counted how many of those variants a pipeline had captured. Annotation of all transcripts in a complex eukaryotic genome is a very difficult task, and even for the human genome, current annotation is not yet stable (Amaral et al., 2023). Therefore, we did not expect automated pipelines to do very well in absolute terms, although their relative performance is nonetheless instructive. We also note that for these six target species, the majority of reference annotations do not have exons that are fully non-coding in the UTRs, which provides an advantage to those methods that do not predict UTRs.

**Figure 3.**
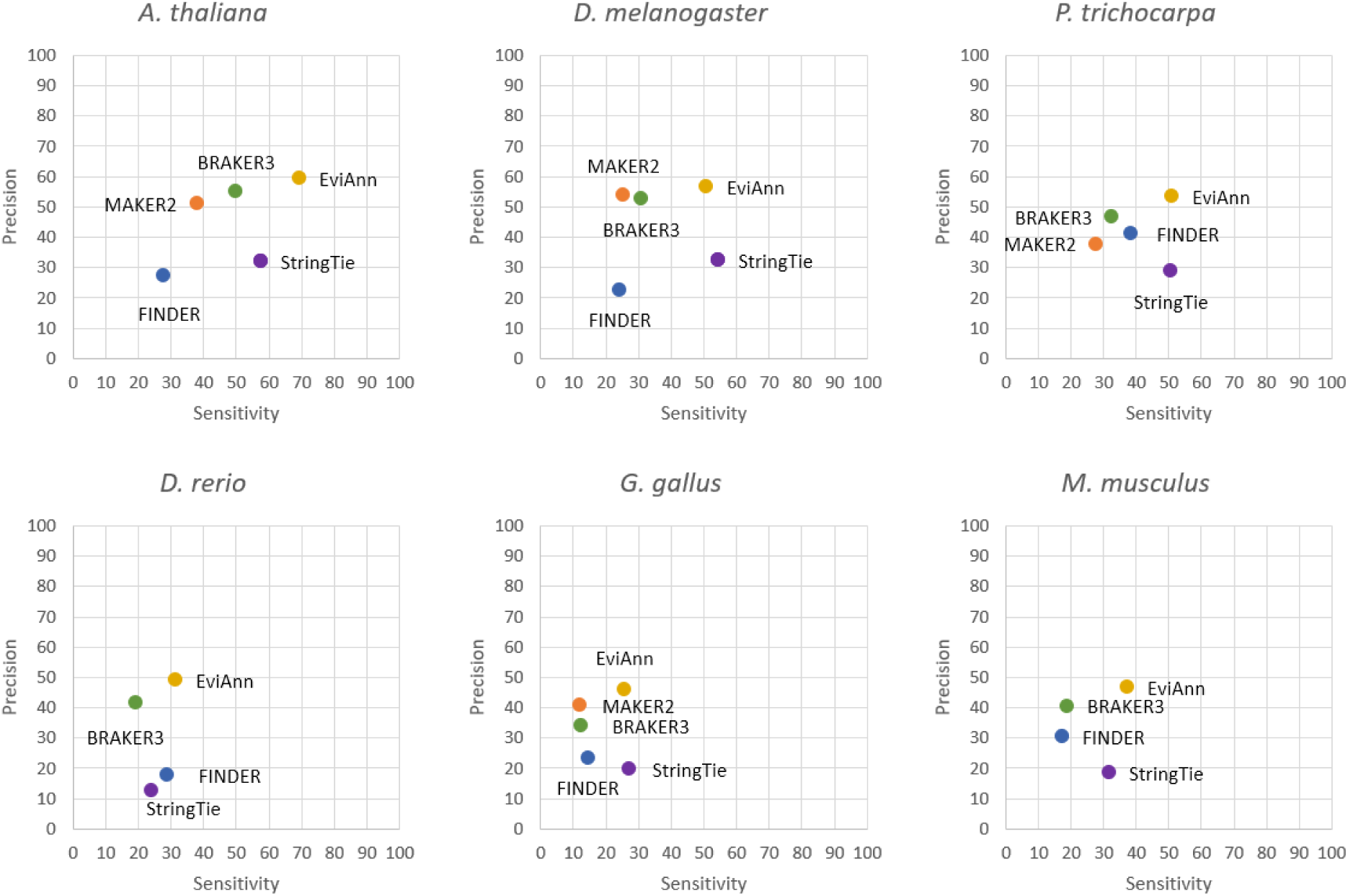
Accuracy of annotations of transcript sequences by EviAnn, MAKER2, BRAKER3, and FINDER in six species. A transcript was considered correct if all introns precisely matched the reference annotation and if both the transcription start and end sites were within 100bp of the reference values. MAKER2 results are not shown for *D. rerio* and *M. musculus* because it failed to complete after one month on a 24-core Intel Xeon Gold server. Note that BRAKER3’s performance in this evaluation was relatively lower because it does not predict long non-coding RNAs or UTRs.

Figure 3 shows that EviAnn consistently achieved the highest sensitivity and precision at the level of transcripts. We note that all pipelines demonstrated relatively low transcript-level sensitivity on zebrafish, chicken, and mouse, likely because of the relatively small amount of RNA-seq data used. For those species, the reference annotation often contains multiple transcripts per gene, which are difficult to capture without large numbers of RNA-seq experiments.

### Comparison of *A. thaliana* annotation to reference annotations of varying confidence levels

For *A.thaliana* we took advantage of the star-rating system used by TAIR10 to do a finer-grained evaluation, in which we separately computed the sensitivity of EviAnn on transcripts with ratings ranging from five stars (highest confidence) down to one star (Table 1). EviAnn’s sensitivity was consistently higher as the confidence rating increased, with just one exception. Curiously, at the transcript level, sensitivity dropped slightly for 5-star transcripts versus those with 4-stars and above. When we investigated, we determined that this drop was due to many 5-star transcripts being annotated without UTRs, which incorrectly penalized EviAnn for including UTR sequences in its predictions for those genes.

**Table 1.** Sensitivity of EviAnn’s annotations for *A. thaliana*, compared to TAIR10 transcripts at various confidence levels.

|  |  |  |  |  |
| --- | --- | --- | --- | --- |
| Transcript rating | Unique transcripts in the reference | Sensitivity vs. annotated transcripts (%) | Sensitivity vs. annotated proteins (CDSs) (%) | Sensitivity vs. annotated genes (%) |
| 5-stars | 8,950 | 75 | 85.8 | 95.8 |
| 4-stars or more | 27,251 | 76.1 | 82.3 | 95.1 |
| 3-stars or more | 30,075 | 74.6 | 80.7 | 94.4 |
| 2-stars or more | 32,737 | 70.1 | 75.6 | 90.3 |
| All TAIR10 transcripts | 35,145 | 69.3 | 74.2 | 87.7 |

### Comparison of BUSCO scores for different annotations

In addition to using reference genome annotations to measure accuracy, we used BUSCO (v5.8.3) (Seppey et al., 2019) in an effort to measure annotation completeness. BUSCO uses lists of single-copy protein-coding genes for a variety of phylogenetic groupings, and the BUSCO software evaluates how many of these genes are completely contained in a genome’s annotation; i.e., for each protein, it determines whether 100% of that protein’s sequence is covered by an annotated gene. As shown in Table 2, EviAnn and BRAKER3 both perform very well using this metric on all six species, with both programs identifying ∼99% of the BUSCO proteins for Arabidopsis, fruit fly, and poplar. For zebrafish, chicken, and mouse, EviAnn captured 91-97% of the BUSCO genes and outperformed BRAKER3 by 4-7%.

**Table 2.**
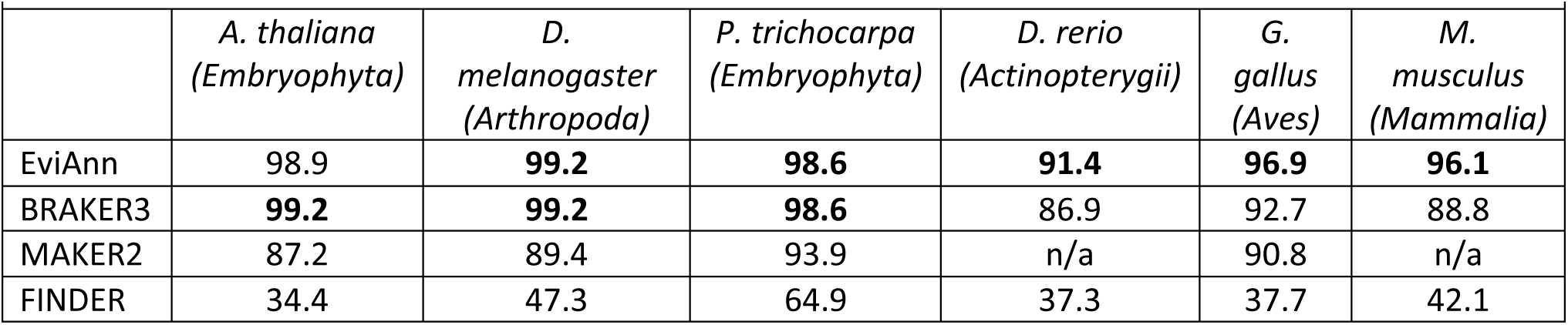
Percentage of complete BUSCO protein sequences found in the annotated proteins produced by EviAnn, BRAKER3, MAKER2, and FINDER for six different species. Columns show the identity of the species as well as the BUSCO protein set used, identified by its phylum name in parentheses. The best results are shown in boldface.

|  | *A. thaliana*<br>(Embryophyta) | *D. melanogaster*<br>(Arthropoda) | *P. trichocarpa*<br>(Embryophyta) | *D. rerio*<br>(Actinopterygii) | *G. gallus*<br>(Aves) | *M. musculus*<br>(Mammalia) |
| --- | --- | --- | --- | --- | --- | --- |
| EviAnn | 98.9 | **99.2** | **98.6** | **91.4** | **96.9** | **96.1** |
| BRAKER3 | **99.2** | **99.2** | **98.6** | 86.9 | 92.7 | 88.8 |
| MAKER2 | 87.2 | 89.4 | 93.9 | n/a | 90.8 | n/a |
| FINDER | 34.4 | 47.3 | 64.9 | 37.3 | 37.7 | 42.1 |

### Annotations of non-canonical splice junctions

Although the vast majority of introns are flanked by the dinucleotides GT/AG in most plant and animal species, many species utilize non-canonical splice sites in a small fraction of their genes. Annotation pipelines vary in their ability to annotate these non-canonical splice sites properly, in part because *ab initio* gene finders rely heavily on the presence of GT/AG signals. We therefore examined the number of non-canonical splice sites in the reference annotation for the six species in this study, and computed how many of these were correctly annotated (according to the reference) by the respective pipelines. As shown in Table 3, although no pipeline consistently predicts these non-canonical splice junctions very well, EviAnn captures far more of the non-canonical splice sites for all six organisms compared to the other pipelines, up to three times as many (in fruit fly) as the next best method.

**Table 3.** Number of non-canonical (non-GT or non-AG) splice sites captured by EviAnn, BRAKER3, MAKER2, and FINDER based on the reference annotations of six plant and animal species. The row labeled Reference shows the total number of non-canonical sites in the reference annotation.

|  | *A. thaliana* | *D. melanogaster* | *P. trichocarpa* | *D. rerio* | *G. gallus* | *M. musculus* |
| --- | --- | --- | --- | --- | --- | --- |
| Reference | 2133 | 6027 | 15628 | 4938 | 32056 | 16925 |
| EviAnn | **1173** | **706** | **2917** | **1898** | **2006** | **1595** |
| BRAKER3 | 864 | 239 | 1724 | 652 | 830 | 844 |
| MAKER2 | 619 | 140 | 1215 | n/a | 790 | n/a |
| FINDER | 486 | 173 | 2203 | 391 | 1090 | 856 |

### Accounting for UTR regions in evaluations

It is important to note that BRAKER3 relies heavily on the AUGUSTUS and GeneMark *ab initio* gene finders, neither of which can annotate UTR regions of transcripts. Nearly all transcripts in the species included in this study begin and end with untranslated regions (UTRs), and for protein coding sequences, the start codon is usually (but not always) found in the first exon, and the stop codon in the last exon. In many cases, though, transcripts begin or end with exons that are entirely noncoding. Genome annotation files normally describe coding regions using CDS features, which are separate from exon features, although their positions overlap. For example, a 4-exon gene might have a coding region starting in exon 2 and ending in exon 4, in which case the annotation will include 4 “exon” features and 3 “CDS” features, where the first CDS will begin somewhere in the middle of exon 2. Although the BRAKER3 system reports only “exon” features in its output, those features correspond only to the protein-coding portions of exons; i.e., they match the CDS features in standard annotation databases. In a recent publication describing BRAKER3 (Gabriel et al., 2024) the authors reported transcript-level accuracy based only on the coding sequences, which incorrectly inflated accuracies because most of the BRAKER3-predicted transcripts were entirely missing their UTRs. In our comparisons here, we have corrected this reporting error, and in Figure 3 we evaluate accuracy on transcripts separately from accuracy on CDS regions, which are evaluated in Figure 2. We further note that BRAKER3 also lacks the ability to annotate long non-coding RNA genes, which number in the thousands for many plants and animals. EviAnn’s ability to use RNA-seq data to annotate long non-coding RNAs contributes to its higher sensitivity on transcripts.

When evaluating protein-coding transcripts predicted by EviAnn, we can break them into three categories: transcripts with RNA-seq evidence, transcripts with protein alignment evidence, and transcripts with both types of evidence. Table 4 provides a breakdown of the numbers of each transcript type in EviAnn’s annotation for the six species used in our experiments. In most cases, both transcript and protein evidence were used by EviAnn, and the number of transcript-only gene loci (genes whose only evidence came from aligning transcripts from a related species to the target genome) was relatively small. A higher number of transcripts with protein-only evidence suggests that the RNA-seq data was not sufficiently deep to capture all distinct proteins for each species. EviAnn includes a label in its output file that indicates the type of evidence used for each annotated transcript, as well as the IDs of transcripts and proteins used as evidence for each annotation.

**Table 4.** Breakdown of the number of protein-coding transcripts annotated with different evidence types for six plant and animal genomes. Evidence support includes “complete” (RNA-seq or related species’ transcripts plus aligned proteins), “transcript only” (transcripts from related species aligned to the genome, but no protein evidence), and “protein only” (protein sequences from related species aligned to the genome, but no transcript evidence).

| Genome | Number of transcripts or coding sequences |  |  |
| --- | --- | --- | --- |
|  | Evidence: complete<br>(UTRs annotated) | Evidence: transcript<br>only (UTRs annotated) | Evidence: protein only<br>(UTRs not annotated) |
| *A. thaliana* | 30183 | 203 | 10574 |
| *D. melanogaster* | 22346 | 494 | 3439 |
| *P. trichocarpa* | 41877 | 374 | 13427 |
| *D. rerio* | 21501 | 356 | 20312 |
| *G. gallus* | 37319 | 432 | 8473 |
| *M. musculus* | 28233 | 359 | 17689 |

### Comparison with more distant protein evidence

When annotating a genome for which there are relatively few protein sequences available from close relatives, EviAnn can use proteins from more-distant species as evidence. To illustrate its performance in this context, we annotated *A. thaliana* using the RNA-seq reads that were used to produce Figures 1–3, but this time we used two different sets of more distant proteins. For the first experiment we used all 305,854 proteins (from RefSeq release 231) from the *Brassiceae* tribe (tribe is a taxonomic rank between family and genus), excluding proteins from the *Camelineae* tribe to which *A. thaliana* belongs. For the second experiment we used as protein evidence 929,959 proteins from the orders *Caryophyllales and Rosales* (*A. thaliana* belongs to the order *Brassicales*). We ran EviAnn, BRAKER3, and MAKER2 using these two sets of proteins from the more distantly related species, and compared the results, using the TAIR10 annotation as a reference. We excluded FINDER from this comparison because it was substantially inferior in all previous experiments. As shown in Figure 4(a), when using the more distant protein set, BRAKER3 has slightly higher sensitivity than EviAnn at the CDS level, but EviAnn was better on both sensitivity and precision at the transcript level. At the gene level, both programs performed similarly. When we used even more distant proteins shown in Figure 4(b), BRAKER3 again showed the best performance at the CDS level, while EviAnn still led at the transcript level. We also included an additional benchmark for these two experiments to showcase EviAnn’s ability to combine a set of externally annotated CDS records with its own annotation (“-c” option). This allowed us to combine the annotations from both BRAKER3 and EviAnn, which is shown on Figure 4 as “EviAnn+BRAKER3” results. This mode showed improved sensitivity at the cost of a small loss in precision.

**Figure 4.**
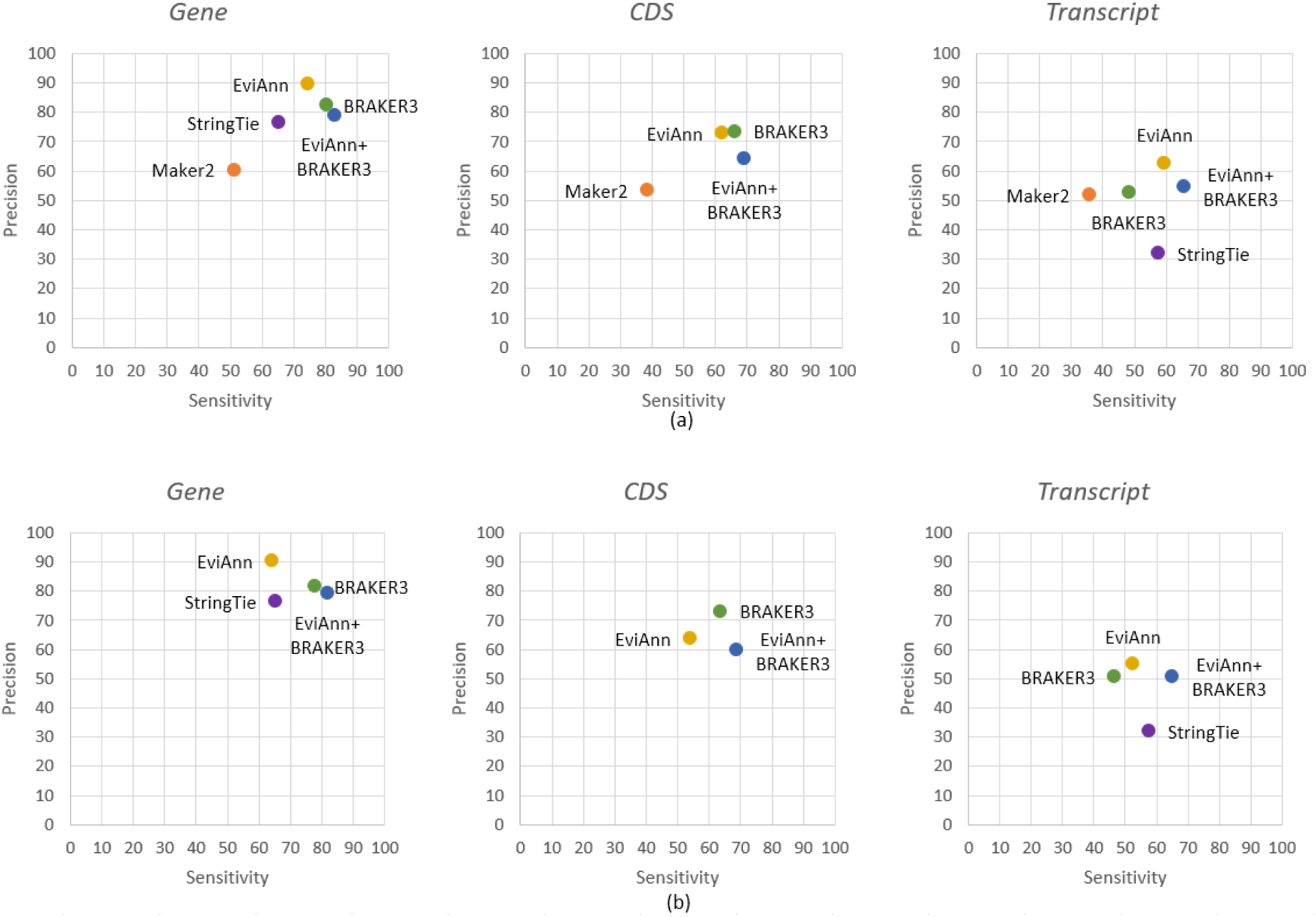
Annotation comparisons for *A. thaliana* using: (a) proteins from a different taxonomic tribe, *Brassiceae*, as input to all pipelines, and (b) using proteins from different taxonomic orders *Caryophyllales* and *Rosales* (*A. thaliana* belongs to order *Brassicales*). Accuracies are shown at the gene, CDS, and transcript levels, defined as in Figures 1–3. MAKER2 failed to run on the set of proteins used for (b) and therefore its results are not shown. Note that BRAKER3’s performance on the transcript level evaluations was relatively lower because it does not predict UTRs.

### How much RNA-seq data is needed?

Because the EviAnn software is transcript-centric, its results should improve with more complete transcript evidence. As we show above, for organisms with smaller genomes (100-200 Mbp), as few as 5-10 RNA-seq data sets may be sufficient to produce high quality annotation. For organisms such as mammals or plants with large genomes, more RNA-seq data sets will likely improve the system’s performance, although it will run robustly with fewer. To provide better guidance on this question, we ran EviAnn on the mouse genome using the same set of proteins as for the previous experiments, but increasing the number of RNA-seq experiments from 6 to 60, with results shown in Figure 5. (SRA IDs for the RNA-seq data sets are listed in Supplementary Table S4.) The standard of truth for these experiments was the NCBI RefSeq annotation for mouse. As expected, EviAnn’s sensitivity for coding sequences and transcripts gradually increased with more data, although it levelled off above 30 data sets. Because the experiments assumed (conservatively) that any prediction not matching RefSeq was a false positive, precision slightly declined with more data, even though some of the additional gene predictions made by EviAnn might be true positives.

**Figure 5.**
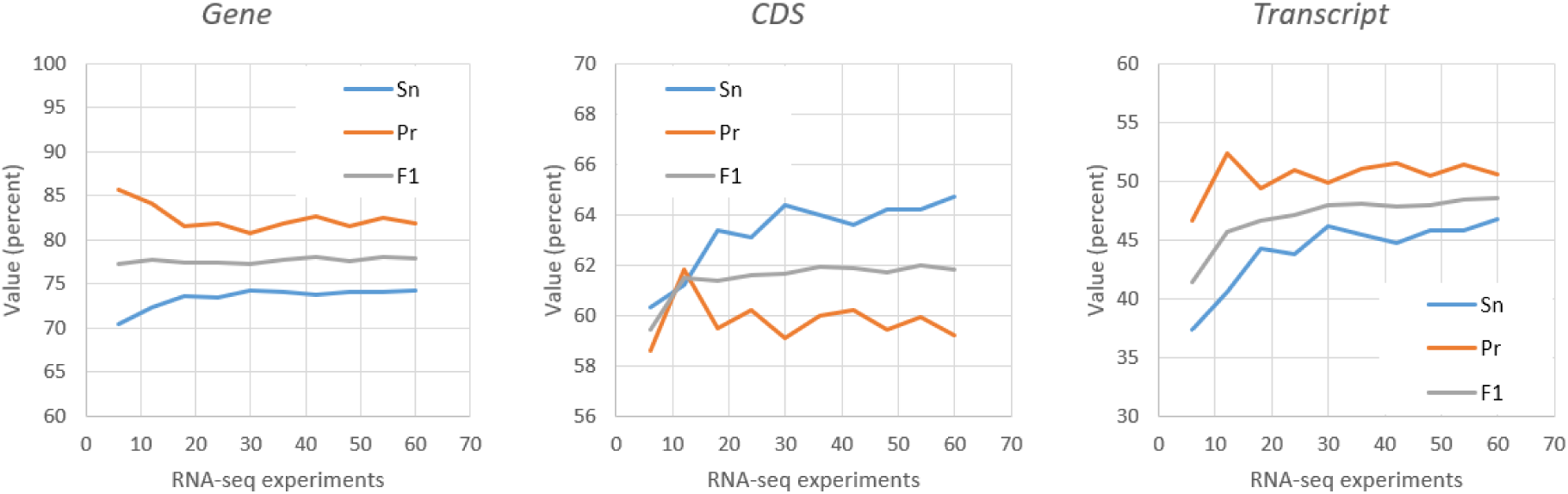
Sensitivity and precision of EviAnn for annotating the mouse (*M. musculus*) genome with a range of input RNA-seq data sets varying from 6 to 60. The individual points are labeled with the number of RNA-seq experiments used. The input protein set was the same for all experiments. The reference annotation used to define true genes, CDSs, and transcripts was the mouse annotation from RefSeq release 231.

### Compute performance

We measured run times for EviAnn, BRAKER3, MAKER2, and FINDER on a Linux server equipped with two 12-core Intel Xeon Gold 6248R CPUs and 1 TB of RAM. We ran all programs using 24 cores. Table 5 lists the timings for all programs on each of the six test genomes, excluding the step of aligning RNA-seq reads to the genome, which was shared by all programs. EviAnn ran far faster on all genomes, with run times ranging from 14 times faster (on mouse, as compared to FINDER) to hundreds of times faster (as compared to MAKER2). EviAnn’s core algorithms, which produce the annotation by examining transcripts and proteins, are very fast, taking up less than 1% of the total run time. The most time-consuming step in EviAnn’s execution is alignment of proteins to the genome, which is done efficiently with miniprot.

**Table 5.** Wall-clock time in hours used by the annotation pipelines EviAnn, BRAKER3, FINDER, and MAKER2 on six different genomes. Timings exclude the time spent on aligning RNA-seq reads to the genome. The fastest result is shown in bold. MAKER2 results for *D. rerio* and *M. musculus* are not shown because the program did not finish after more than one month of run time.

|  |  |  |  |  |  |  |
| --- | --- | --- | --- | --- | --- | --- |
|  | *A. thaliana* | *D. melanogaster* | *D. rerio* | *P. trichocarpa* | *G. gallus* | *M. musculus* |
| EviAnn | **0.3** | **0.3** | **1.0** | **0.4** | **0.4** | **0.8** |
| BRAKER3 | 13.2 | 7.3 | 29 | 10.8 | 72 | 51 |
| FINDER | 6.8 | 8.4 | 63 | 9.5 | 38 | 11 |
| MAKER2 | 168.0 | 34.0 | N/A | 81.0 | 148 | N/A |

## Methods

### Input data

The overall annotation pipeline for EviAnn is illustrated in Figure 5. As input, EviAnn uses the genome to be annotated, RNA-seq data or transcripts from one or more closely related species, and alignments of protein sequences to the genomic DNA. RNA-seq data must be grouped in such a way that each input file (or pair of files in the case of paired-end sequencing) represents a single RNA sequencing run; e.g. a single condition for a single tissue. EviAnn supports sequencing data from Illumina RNA-seq, PacBio IsoSeq, and Oxford Nanopore direct RNA/cDNA, as well as mixed input containing both short-and long-read sequences. EviAnn can also use transcripts from different but closely related species, which are supplied as separate files. EviAnn uses HISAT2 to align short-read RNA-seq data to the genome, and it uses minimap2 to align long read RNA-seq data and/or transcripts from related species. EviAnn uses the resulting spliced alignments of reads and transcripts as input to the transcriptome assembly program StringTie2 (Kovaka et al., 2019). Alignments from short-read RNA-seq experiments are assembled separately from long read data. For RNA-seq experiments, EviAnn keeps any transcripts with a TPM > 0.25 that are present in at least 10% of the input experiments, or that have a TPM expression level of at least 100 in any one sample. Note that if fewer than 10 data sets are provided, transcripts need only be present in one sample with TPM > 0.25. After assembling all RNA-seq data, EviAnn uses gffcompare (Pertea and Pertea, 2020) to eliminate redundant and fragmented transcripts and to merge all transcript assemblies into a single transcript set. After merging, EviAnn assigns to each transcript a TPM value equal to the maximum TPM across all input samples, and encodes this value along with the number of samples in the transcript name.

### Merging transcript and protein alignments

EviAnn’s use of protein sequences from related species works as follows. It first aligns proteins to genomic DNA using miniprot (Li et al, 2023), and then it runs gffcompare with the protein alignment file, and uses those results to assign CDS coordinates to each assembled transcript. EviAnn then examines all transcripts that share at least one intron junction (or that overlap, in the case of intronless transcripts) with the protein alignment. EviAnn may keep more than one transcript corresponding to the same protein if they have different UTR regions. If multiple transcripts contain the same protein at the same locus, EviAnn computes a transcript “reliability” score *R* for all such transcripts. *R* is computed as *N*log_2_(2+T),* where *N* is the number of RNA-seq experiments in which the transcript was observed, and *T* is the TPM value. This score gives greater weight to transcripts that are expressed in multiple tissues and at higher levels. For each gene, EviAnn computes a maximum score *R_max_*, and keeps only those transcripts whose *R* score is at least √*R_max_*, a threshold that was chosen empirically. EviAnn labels transcripts that do not have any corresponding intron chain matches to a protein as “potentially noncoding” transcripts.

At the completion of this stage, we have three categories of genes:

1. Genes with transcripts that have a complete intron chain match to a CDS from an aligned protein (“complete”);
2. Genes with transcripts where none of the transcripts have a corresponding protein alignment (“transcript-only”); and
3. Genes whose coding sequences are annotated based only on protein alignments, with no RNA-seq support and no UTR regions (“protein-only”).

Next, EviAnn constructs a Markov chain model to screen out and remove transcripts with low-quality splice sites. Note that while some *ab initio* gene finders use Markov chains to identify splice sites, in EviAnn we employ them only as a filtering step on transcripts that were constructed by other methods, as shown in Figure 6. We model donor and acceptor splice sites separately, as follows. First, we label as *reliable* all complete transcripts that do not have introns in the UTR regions, and whose CDSs contain both start and stop codons with no in-frame stops. (Note that using this definition, >99.8% of all introns in reliable transcripts match RefSeq annotations exactly.) We then use all unique introns in reliable transcripts to compute Markov chain model weights for donor and acceptor splice sites for the target genome. For the donor site, EviAnn builds a Markov chain spanning 16 positions, with the consensus GT (the beginning of the intron) at positions 4-5. For acceptor sites, it creates a 30-bp Markov chain where the consensus AG is at positions 26-27. If the data contains fewer than 1024 splice sites, EviAnn uses a very simple 0^th^-order chain (equivalent to a position weight matrix, or PWM), with 4 probabilities at each position. If the data contains 1024–4096 splice sites, we use a 1^st^-order Markov chain, where the probability of each base in the chain is dependent on one position to its immediate left, with 16 probabilities at each position. If the data contains >4096 sites, we use a 2^nd^-order Markov chain, with 64 probabilities per position. EviAnn uses its Markov chain to score every splice junction in the preliminary annotation, and scores each intron as the sum of Markov chain scores for donor and acceptor sites. We then assign a score to each transcript equal to the minimum intron score over all introns in the transcript. Finally, EviAnn eliminates transcripts and aligned CDSs where the minimum intron score is below a pre-specified threshold, which is set to 4 based on the test data sets used in this study.

**Figure 6.**
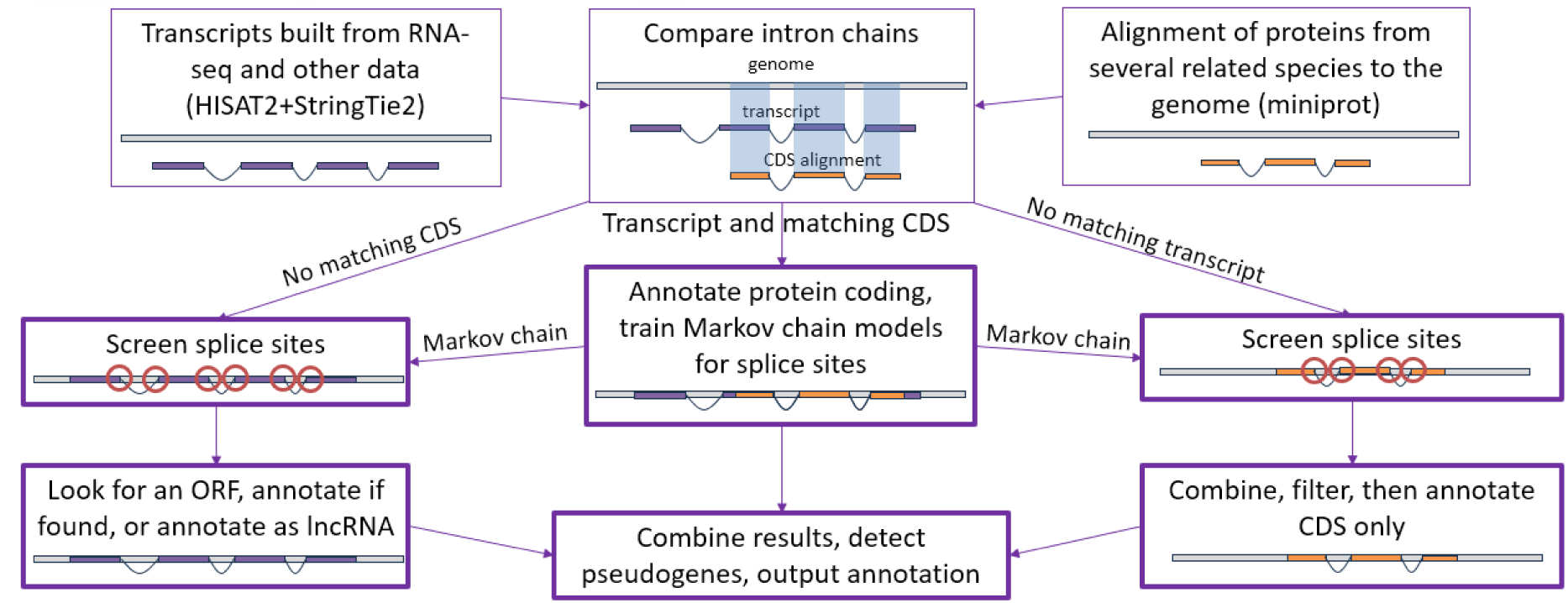
A simplified diagram of the EviAnn genome annotation pipeline.

Sometimes one or more junctions in an intron chain for a transcript might disagree with its best matching CDS. In the case of disagreements such as this, EviAnn will construct the longest possible transcript model with the largest number of exons derived from the transcript and the CDS alignments, as shown in Extended Data Figure 1. For each transcript, EviAnn first assumes that the intron junctions derived from the transcript are correct, and it infers putative start and end coordinates of the open reading frame (ORF) from the corresponding start and stop of the CDS (derived from the protein alignment). EviAnn then verifies that the ORF has no in-frame stop codons, its length is divisible by 3, it starts with a start codon (ATG), and ends with a stop codon (TAA, TAG, or TGA). We observed that most ORFs have correct start and stop codons and no in-frame stops at this step.

If an ORF has an in-frame stop or its length is not divisible by 3, EviAnn uses TransDecoder (Haas, 2023) to look for a complete ORF that has an amino-acid match to the original aligned protein. If TransDecoder is unable to find an ORF that satisfies these criteria, the transcript is discarded. If a start codon is missing, EviAnn searches for the furthest start codon in the 5’ direction, starting with the implied beginning of the CDS, and if not found, it searches in the 3’ direction. If a start codon is found and the stop codon is missing, EviAnn looks along the transcript sequence for a first stop codon downstream from the start codon. No frameshifts are allowed when searching for start or stop codons. If EviAnn fails to find a complete ORF, and there was a disagreement between the intron chains for the transcript and its best matching CDS, it discards the transcript and attempts to find a complete ORF in the aligned CDS alone, repeating the above process for the CDS.

For transcript-only gene loci; i.e., those with no protein sequence matches, EviAnn considers whether these might contain novel protein-coding genes or instead represent noncoding RNA genes. EviAnn uses TransDecoder to find complete ORFs in these transcripts, and if it finds an ORF spanning > 75% of the transcript, that gene is annotated as protein-coding, and all other transcripts at that locus that do not contain a CDS are discarded. If none of the transcripts at a locus have a valid ORF that satisfies the above conditions, EviAnn declares the locus to be non-coding and annotates all transcripts at that locus as long non-coding RNAs.

Protein-only annotations contain genes that EviAnn identified solely on protein alignments, without either RNA-seq or transcript alignment support. For this class of genes, we only use protein alignments that yield a complete open reading frame in the target genome, whose alignment spans the entire source protein without any frameshifts in the coding region. Because these annotations lack transcript evidence, the annotated transcripts will lack UTRs. EviAnn annotates UTRs only when it has transcripts obtained from StringTie assemblies of RNA sequencing data, or when it can align previously-annotated transcripts taken from very closely related species.

Note that after creating its initial annotation, EviAnn looks for UTR regions that overlap the CDS regions of other genes. Transcripts with such UTRs might represent “readthrough” transcripts that include parts of two neighboring genes, but EviAnn does not include these in its output. If it finds overlapping UTRs, EviAnn splits the overlapping transcripts to remove the overlap.

### UTR annotations

UTRs are only annotated for protein coding transcripts obtained from RNA sequencing evidence such as transcript assemblies by StringTie or alignments of transcripts from closely related species. For transcripts derived from RNA sequencing data, EviAnn relies on StringTie transcript assemblies to determine the 5’ TSS and 3’ TTS boundaries. StringTie determines these using a majority-weighted method which is internal to the StringTie algorithm; e.g., it uses the coordinates where most of the reads begin and end to determine the UTR boundaries. If a multi-exon transcript with the same intron chain is present in several RNA-seq samples, we compute 5’ TSS and 3’ TTS locations averaging the transcript beginning and end positions over all samples.

### Processed pseudogene detection and optional functional annotation

Processed pseudogenes are copies of spliced mRNAs (i.e., transcripts with introns removed) that have been incorporated into the genome sequence. While some of these genes may have a valid ORF, they are generally non-functional and should not be annotated as protein-coding genes. At the last step of the EviAnn pipeline, it uses blastp (Camacho et al., 2009) to align all proteins from intronless transcripts to all proteins coded by multi-exon transcripts. If an intronless protein aligns with >90% similarity over >90% of its length to a multi-exon protein, EviAnn reports it as a pseudogene, and it does not output CDS records for such transcripts.

Optionally, if the user provides the -f switch to EviAnn, it aligns all annotated proteins to the UniProt-SwissProt protein database with blastp. It then uses the name of the best matching protein to assign a name to the newly annotated protein, with the note “Similar to” prepended.

### Performing the evaluations

We ran EviAnn, MAKER2, BRAKER3, and FINDER on six plant and animal species: *Arabidopsis thaliana*, *Drosophila melanogaster*, *Populus trichocarpa* (poplar tree), *Danio rerio* (zebrafish), *Gallus gallus* (chicken), and *Mus musculus* (mouse). We used repeat-masked genomes downloaded from the NCBI RefSeq database; accession numbers are shown in Supplementary Table S3. Even though EviAnn does not require the input genome to be repeat-masked, the other software pipelines do.

For RNA sequencing and protein sequence inputs to each of the annotation systems, we used the “close relatives” data sets from Gabriel et al. (2024). These data sets contain RNA sequencing data available from NCBI plus proteins from several closely-related species. Accession numbers for each data set are listed in Supplementary Tables S1 and S2.

To process RNA-seq reads, EviAnn uses HISAT2 (Kim et al., 2019) to align the reads to the target genome. We provided the same alignments to BRAKER3 and FINDER. Note that EviAnn and BRAKER3 use StringTie2 internally to assemble transcripts from the aligned RNA-seq reads. As part of our transcript-level evaluations, we included the StringTie-assembled transcripts, which we merged using gffcompare, to show that the annotation pipelines provided further improvements beyond what StringTie2 alone produced.

MAKER2 is unable to use aligned RNA-seq data directly and instead it uses transcripts as “EST” evidence for its initial evidence-based annotation pass. We provided the StringTie-assembled transcripts as “EST” evidence to MAKER2. The MAKER2 pipeline also requires manual intervention to train an external gene finder; in our experiments, we trained SNAP (Korf, 2004) following the MAKER2 protocol (Campbell et al., 2014). For convenience and for better performance in terms of both speed and accuracy, we developed an automated script to run MAKER2 in parallel on a single multi-core computer. This script, which we called EZMAKER, is included in the EviAnn distribution package as `ez_maker.sh`, and serves as a wrapper to run two-pass annotation (using the -d switch, as we did here) with MAKER2, incorporating the SNAP *ab initio* gene finding step. This wrapper makes it easier to reproduce the results described here, and in addition it makes MAKER2 faster and easier to run on a single multi-core server. We list the command lines to generate all annotations in the Supplementary Materials.

### Metrics of annotation quality

We used gffcompare (Pertea and Pertea, 2020) to compare the automatically generated annotations to the protein-coding and lncRNA genes in the NCBI RefSeq annotations of all species except *A. thaliana*, for which we used the TAIR10 annotation (Lamesch et al., 2012). Note that our evaluation specifically assessed the annotated transcripts and CDSs against the complete RefSeq transcripts. Thus, if an annotated transcript was missing an exon in either the 5’ or 3’ UTR regions, it was not considered a match. We did, however, allow mismatches of up to 100bp at both the transcriptional start and the transcription termination sites. We evaluated accuracy on protein-coding sequences (CDSs) separately.

We assessed the various annotations using sensitivity and precision, computed as:

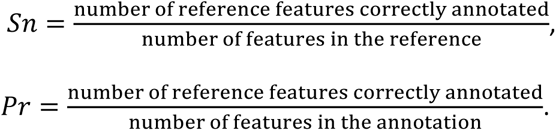

We also computed F1 scores as:

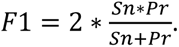

Because F1 is a composite of sensitivity and precision and thus somewhat redundant, we report F1 results in Supplementary Tables S5-S7. We report total counts of genes, transcripts and unique proteins annotated by all pipelines in Supplementary Table S8. We computed these statistics for three feature types: genes (or gene loci), CDSs, and transcripts. A gene was considered correctly annotated if at least one transcript or CDS at the gene locus matched the reference annotation, as determined by either a perfect intron chain match (for multi-exon transcripts) or exon coordinate overlap (for intronless transcripts). A CDS match was considered valid if and only if the start and end of translation and all introns within the coding region matched exactly. Finally, a transcript was considered valid if all intron coordinates matched exactly and transcription start and end sites were both within 100 bp of the reference sites. For intronless transcripts, we required that at least 80% of the transcript had to overlap the reference transcript. Note that in Figures 1-3, we do not show MAKER2 results for *D. rerio* and *M. musculus* because MAKER2 failed to complete the annotation in one month on a 24-core Intel Xeon Gold server.

## Discussion

EviAnn uses a combination of transcription evidence and protein sequence alignments to produce automated, high-quality eukaryotic genome annotation, outperforming most existing packages in almost all measures of accuracy. EviAnn’s ability to use transcripts and proteins from close relatives enables the annotation of novel genomes even when expression data is unavailable. With the ever-increasing number of well-assembled and well-annotated genomes available from public databases, we expect EviAnn to become even more useful over time.

EviAnn was designed with the goals of transparency and efficiency in mind. When EviAnn is provided assembled transcripts as input, any transcript that aligns well with the DNA of the target genome, and that has a valid open reading frame whose translation matches a known protein, will be included in the output. The system’s output includes labels that describe the evidence that was used to support each transcript, allowing users to easily see whether a particular transcript is supported by expression data, protein alignments, or both.

EviAnn does not require the input genome to be masked for repeats, whereas all competing annotation pipelines that utilize *ab initio* gene finders do. Repeat masking is a computationally intensive process, and eliminating the need for a repeat-masked genome saves both time and effort in genome annotation projects.

Finally, we note that if a well-annotated genome from the same or a very closely related species is available, annotation liftover can be used to annotate the target genome by mapping genes and other features directly between the two genomes. We do not recommend using EviAnn as the primary tool for this purpose, since other tools such as Liftoff (Shumate and Salzberg, 2020) or LiftOn (Chao et al., 2025) were specifically developed for annotation liftover and are better suited to this task.

## Acknowledgements

This work was supported in part by NSF grant IOS-2432298, and by NIH grants R01-HG006677, R35-GM130151, and R35-GM156470.

## Author Contributions Statement

**Aleksey V. Zimin:** Conceptualization, Methodology, Software, Validation, Writing - Original Draft, Writing - Review & Editing, Visualization, Project administration, Funding acquisition.

**Daniela Puiu:** Validation, Writing - Review & Editing, Visualization.

**Mihaela Pertea:** Conceptualization, Methodology, Writing - Review & Editing.

**James A. Yorke:** Conceptualization, Methodology.

**Steven L. Salzberg:** Conceptualization, Methodology, Validation, Writing - Review & Editing, Project administration, Funding acquisition.

## Figure Legends (Extended Data)

**Extended data Figure 1.**
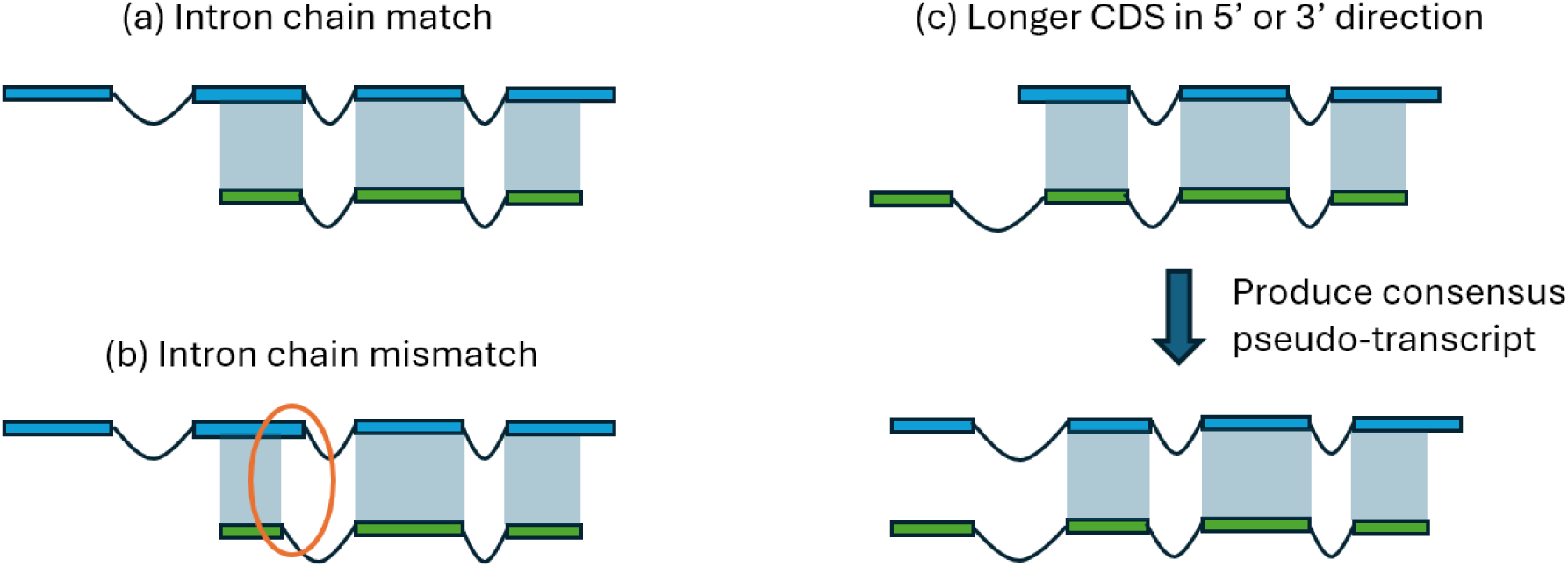
Resolution of conflicts between intron chains of aligned coding sequences (CDSs) from related species’ proteins (green) and transcripts assembled from the RNA-seq data (blue). In all cases, EviAnn uses the frame specified by the start of the aligned CDS and only annotates a transcript/CDS pair if it can locate a complete ORF on the transcript. (a) If there are no conflicts in the intron chain, use the transcript/CDS pair as is, and look for a complete ORF. (b) If the intron chain fails to match, trust the transcript intron chain (blue), look for a complete ORF on the transcript, and annotate if found. If no complete ORF is found, delete the transcript and use the CDS if it contains a complete ORF. (c) With a partial intron chain match, look for the first matching splice junction (arrow) and produce a consensus pseudo-transcript using exons from the transcript and CDS. Not pictured here: if the transcript is contained in the CDS, simply discard the transcript and use the exons from the CDS.

## Data Availability

NCBI accession numbers for all data sets used in this study are provided in Supplementary Tables S1-S4. The 305,854 proteins (from RefSeq release 231) from the *Brassiceae* tribe can be found at NCBI under taxonomy ID 981071. The 929,959 proteins from the orders *Caryophyllales and Rosales* from RefSeq release 231 can be found at NCBI under taxonomy IDs 3524 and 3744.

## Code Availability

Eviann’s source code is freely available under the GPL-3.0 open-source license from github at https://github.com/alekseyzimin/EviAnn_release/, and though Bioconda as “eviann”.

