## Supplementary Materials for "Efficient evidence-based genome annotation with EviAnn"

### Supplementary Materials for the manuscript "Efficient evidence-based genome annotation with EviAnn"

**Data used for evaluations.** We used the same data sets for our comparisons as were used in the "closest relatives" data set in (Gabriel *et al.*, 2024). The species whose proteins were used as the "closest relatives" set are listed in Supplementary Table 1. Supplementary Table 2 lists the NCBI SRA accession numbers for the RNA-seq data used for annotation experiments. Supplementary Table 3 lists the NCBI RefSeq and GenBank IDs for the reference annotations used for our evaluations.

Supplementary Table 1. NCBI accession numbers for the species whose proteins were used for annotation experiments.

| **Species** | **Proteins NCBI accession** |
|---|---|
| *Arabidopsis thaliana* | |
| *Arabidopsis lyrata subsp. lyrata* | GCF_000004255.2 |
| *Arabidopsis thaliana x Arabidopsis arenosa* | GCA_019202795.1 |
| *Camelina sativa* | GCF_000633955.1 |
| *Arabidopsis suecica* | GCA_019202805.1 |
| *Capsella rubella* | GCF_000375325.1 |
| *Danio rerio* | |
| *Cyprinus carpio* | GCF_018340385.1 |
| *Carassius auratus* | GCF_003368295.1 |
| *Puntigrus tetrazona* | GCF_018831695.1 |
| *Sinocyclocheilus rhinocerous* | GCF_001515625.1 |
| *Sinocyclocheilus anshuiensis* | GCF_001515605.1 |
| *Onychostoma macrolepis* | GCA_012432095.1 |
| *Carassius gibelio* | GCF_023724105.1 |
| *Pimephales promelas* | GCF_016745375.1 |
| *Labeo rohita* | GCF_022985175.1 |
| *Megalobrama amblycephala* | GCF_018812025.1 |
| *Sinocyclocheilus grahami* | GCF_001515645.1 |
| *Ctenopharyngodon idella* | GCF_019924925.1 |
| *Drosophila melanogaster* | |
| *Drosophila ananassae* | GCF_017639315.1 |
| *Drosophila grimshawi* | GCF_018153295.1 |
| *Drosophila pseudoobscura* | GCF_009870125.1 |
| *Drosophila virilis* | GCF_003285735.1 |
| *Drosophila willistoni* | GCF_018902025.1 |

| *Gallus gallus* | |
|---|---|
| *Lagopus muta* | GCF_023343835.1 |
| *Tympanuchus pallidicinctus* | GCF_026119805.1 |
| *Lagopus leucura* | GCF_019238085.1 |
| *Centrocercus urophasianus* | GCF_019232065.1 |
| *Centrocercus urophasianus* | GCF_019232065.1 |
| *Coturnix japonica* | GCF_001577835.2 |
| *Meleagris gallopavo* | GCF_000146605.3 |
| *Mus musculus* | |
| *Arvicanthis niloticus* | GCF_011762505.1 |
| *Grammomys surdaster* | GCF_004785775.1 |
| *Mastomys coucha* | GCF_008632895.1 |
| *Mus pahari* | GCF_900095145.1 |
| *Apodemus sylvaticus* | GCF_947179515.1 |
| *Mus caroli* | GCF_900094665.1 |
| *Rattus rattus* | GCF_011064425.1 |
| *Rattus norvegicus* | GCF_015227675.2 |
| *Homo sapiens* | GCF_000001405.40 |
| *Populus trichocarpa* | |
| *Populus tomentosa* | GCA_018804465.1 |
| *Populus euphratica* | GCF_000495115.1 |
| *Populus alba* | GCF_005239225.1 |
| *Populus deltoides* | GCA_015852605.2 |

Supplementary Table 2. RNA-seq data used for annotation experiments.

| **Species** | **SRA IDs** |
|---|---|
| *A. thaliana* | SRR8714016, SRR8759751, SRR4010853, SRR7289569, SRR12547664, SRR12076896 |
| *D. melanogaster* | SRR19416937, SRR19416947, SRR19416944, SRR19446462, SRR19416948 |
| *P. trichocarpa* | SRR3019959, SRR3019957, SRR3019251, SRR3019585, SRR3019304, SRR12671667 |
| *D. rerio* | SRR3179613, ERR1857957, SRR9159941, SRR9159937, ERR958944, SRR8106574 |
| *G. gallus* | SRR5340686, ERR2113192, ERR2113173, SRR5190436, SRR1822373, SRR5437696 |
| *M. musculus* | SRR5197958, SRR3094250, SRR6067921, SRR10115888, ERR3005082, SRR9202226 |

Supplementary Table 3. NCBI IDs for the species whose reference annotations were used for evaluations.

| **Species** | **NCBI ID for the reference assembly and annotation** |
|---|---|
| *D. melanogaster* | GCF_000001215.4 |

| | |
|---|---|
| *P. trichocarpa* | GCF_000002775.5 |
| *D. rerio* | GCF_000002035.6 |
| *G. gallus* | GCF_016699485.2 |
| *M. musculus* | GCF_000001635.27 |

| Supplementary Table 4. SRA IDs for expanded RNA-seq dataset for *M. musculus* (54 additional experiments/runs). |
|---|
| SRR35819776 |
| SRR35819770 |
| SRR35783298 |
| SRR35783299 |
| SRR35810336 |
| SRR35810335 |
| SRR35940547 |
| SRR35948833 |
| SRR35948834 |
| SRR35948835 |
| SRR35895405 |
| SRR35895404 |
| SRR35910694 |
| SRR35962726 |
| SRR35962725 |
| SRR35996889 |
| SRR35996890 |
| SRR35983974 |
| SRR35983973 |
| SRR35996886 |
| SRR35996885 |
| SRR36008043 |
| SRR35991682 |
| SRR36003884 |
| SRR35991719 |
| SRR36014290 |
| SRR36014291 |
| SRR36014169 |
| SRR36003881 |
| SRR31629396 |
| ERR5310867 |
| SRR14125700 |
| SRR14119964 |
| SRR14128259 |
| SRR14130132 |
| SRR14130376 |

| |
|---|
| SRR23884968 |
| SRR23892263 |
| SRR23886717 |
| SRR23886531 |
| SRR23892249 |
| SRR33548324 |
| SRR23886527 |
| SRR31208698 |
| SRR33481121 |
| SRR33572599 |
| SRR33572586 |
| SRR34468024 |
| SRR34999192 |
| SRR34518226 |
| SRR34517977 |
| SRR34986235 |
| SRR34999102 |
| SRR34986654 |

**Numerical values for the evaluation results shown in Figures 1-3.**

Supplementary Table 5. Numerical values for gene locus level evaluations of the four pipelines on the "closest relatives" data sets described in Tables S1-S3. The best result in each row is in boldface.

| Species | | | *A. thaliana* | | |
|---|---|---|---|---|---|
| | **EviAnn** | **FINDER** | **MAKER2** | **BRAKER3** | **StringTie** |
| Sensitivity | **87.7** | 34.8 | 54.4 | 82.8 | 65.3 |
| Precision | **88.3** | 28.7 | 60.1 | 79 | 76.6 |
| F1 score | **88.0** | 31.5 | 57.1 | 80.9 | 70.5 |
| Species | | | *D. melanogaster* | | |
| | **EviAnn** | **FINDER** | **MAKER2** | **BRAKER3** | **StringTie** |
| Sensitivity | 74.5 | 37.9 | 52.6 | **74.8** | 68.6 |
| Precision | **92.7** | 22.8 | 70.9 | 85.1 | 68.4 |
| F1 score | **82.6** | 28.5 | 60.4 | 79.6 | 68.5 |
| Species | | | *P. trichocarpa* | | |
| | **EviAnn** | **FINDER** | **MAKER2** | **BRAKER3** | **StringTie** |
| Sensitivity | 81.6 | 61.3 | 55 | **82.3** | 71.1 |
| Precision | **84.6** | 48.5 | 44 | 74.8 | 69.8 |
| F1 score | **83.1** | 54.2 | 48.9 | 78.4 | 70.4 |
| Species | | | *D. rerio* | | |
| | **EviAnn** | **FINDER** | **MAKER2** | **BRAKER3** | **StringTie** |
| Sensitivity | **53.9** | 32.7 | | 49.2 | 35.7 |
| Precision | **76.7** | 20.7 | | 73.7 | 29.1 |
| F1 score | **63.3** | 25.4 | | 59.0 | 32.1 |
| Species | | | *G. gallus* | | |
| | **EviAnn** | **FINDER** | **MAKER2** | **BRAKER3** | **StringTie** |
| Sensitivity | **57** | 38.7 | 40.4 | 54.9 | 52.4 |

| | | | | | |
|---|---|---|---|---|---|
| Precision | **85.1** | 25.4 | 54.4 | 55.3 | 51.2 |
| F1 score | **68.3** | 30.7 | 46.4 | 55.1 | 51.8 |
| Species | *M. musculus* | | | | |
| | **EviAnn** | **FINDER** | **MAKER2** | **BRAKER3** | **StringTie** |
| Sensitivity | **70.5** | 35.4 | | 60.6 | 48.4 |
| Precision | **85.6** | 34 | | 74.8 | 45.9 |
| F1 score | **77.3** | 34.7 | | 67.0 | 47.1 |

| Supplementary Table 6. Numerical values for CDS level evaluations of the four pipelines on "Closest relatives" data sets described in Tables S1-S3. The best number in each row is in bold. | | | | |
|---|---|---|---|---|
| Species | *A. thaliana* | | | |
| | **EviAnn** | **FINDER** | **MAKER2** | **BRAKER3** |
| Sensitivity | **74.2** | 7.9 | 41.4 | 68.6 |
| Precision | 69 | 34.7 | 53.2 | **72.5** |
| F1 score | **71.5** | 12.9 | 46.6 | 70.5 |
| Species | *D. melanogaster* | | | |
| | **EviAnn** | **FINDER** | **MAKER2** | **BRAKER3** |
| Sensitivity | **66.5** | 14.9 | 39 | 60.8 |
| Precision | 74.5 | 45.8 | 64.2 | **78.1** |
| F1 score | **70.3** | 22.5 | 48.5 | 68.4 |
| Species | *P. trichocarpa* | | | |
| | **EviAnn** | **FINDER** | **MAKER2** | **BRAKER3** |
| Sensitivity | **66.7** | 21.5 | 39.7 | 63.4 |
| Precision | 62.8 | 34.6 | 40.5 | **68.8** |
| F1 score | 64.7 | 26.5 | 40.1 | **66.0** |
| Species | *D. rerio* | | | |
| | **EviAnn** | **FINDER** | **MAKER2** | **BRAKER3** |
| Sensitivity | **48.6** | 26.2 | | 40.2 |
| Precision | 60.9 | 44 | | **67** |
| F1 score | **54.1** | 32.8 | | 50.3 |
| Species | *G. gallus* | | | |
| | **EviAnn** | **FINDER** | **MAKER2** | **BRAKER3** |
| Sensitivity | **44.1** | 7.7 | 19.6 | 34.1 |
| Precision | **58.5** | 26.7 | 39.8 | 53.6 |
| F1 score | **50.3** | 12.0 | 26.3 | 41.7 |
| Species | *M. musculus* | | | |
| | **EviAnn** | **FINDER** | **MAKER2** | **BRAKER3** |
| Sensitivity | **60.3** | 16.6 | | 45.5 |
| Precision | 58.6 | 51.2 | | **66.5** |
| F1 score | **59.4** | 25.1 | | 54.0 |

| Supplementary Table 7. Numerical values for transcript level evaluations of the four pipelines on "closest relatives" data sets described in Tables S1-S3. The best number in each row is in bold. | | | | | |
|---|---|---|---|---|---|
| Species | *A. thaliana* | | | | |
| | **EviAnn** | **FINDER** | **MAKER2** | **BRAKER3** | **StringTie** |

| | | | | | |
|---|---|---|---|---|---|
| Sensitivity | **69.3** | 27.9 | 37.9 | 50 | 57.5 |
| Precision | **59.6** | 27.3 | 51.2 | 55.2 | 31.9 |
| F1 score | **64.1** | 27.6 | 43.6 | 52.5 | 41.0 |
| Species | *D. melanogaster* | | | | |
| | **EviAnn** | **FINDER** | **MAKER2** | **BRAKER3** | **StringTie** |
| Sensitivity | 50.9 | 24.3 | 25.4 | 30.9 | **54.5** |
| Precision | **56.5** | 22.5 | 53.9 | 52.6 | 32.5 |
| F1 score | **53.6** | 23.4 | 34.5 | 38.9 | 40.7 |
| Species | *P. trichocarpa* | | | | |
| | **EviAnn** | **FINDER** | **MAKER2** | **BRAKER3** | **StringTie** |
| Sensitivity | **51.1** | 38.2 | 27.7 | 32.6 | 50.7 |
| Precision | **53.4** | 41.3 | 37.8 | 46.9 | 28.7 |
| F1 score | **52.2** | 39.7 | 32.0 | 38.5 | 36.7 |
| Species | *D. rerio* | | | | |
| | **EviAnn** | **FINDER** | **MAKER2** | **BRAKER3** | **StringTie** |
| Sensitivity | **31.5** | 28.8 | | 19.3 | 24.2 |
| Precision | **49.2** | 17.8 | | 41.8 | 12.7 |
| F1 score | **38.4** | 22.0 | | 26.4 | 16.7 |
| Species | *G. gallus* | | | | |
| | **EviAnn** | **FINDER** | **MAKER2** | **BRAKER3** | **StringTie** |
| Sensitivity | 25.8 | 14.5 | 11.9 | 12.4 | **27** |
| Precision | **46** | 23.5 | 40.9 | 34.2 | 19.6 |
| F1 score | **33.1** | 17.9 | 18.4 | 18.2 | 22.7 |
| Species | *M. musculus* | | | | |
| | **EviAnn** | **FINDER** | **MAKER2** | **BRAKER3** | **StringTie** |
| Sensitivity | **37.3** | 17.3 | | 18.8 | 31.7 |
| Precision | **46.6** | 30.6 | | 40.6 | 18.6 |
| F1 score | **41.4** | 22.1 | | 25.7 | 23.4 |

Supplementary Table 8. Number of genes, transcripts, long non-coding RNAs (lncRNAs), and unique proteins annotated for each species. For Reference counts, we only used the reliable curated features (transcript or gene IDs starting with N) from RefSeq annotations, except for *A. thaliana* for which we used TAIR10 annotation.

| Species | *A. thaliana* | | | | | |
|---|---|---|---|---|---|---|
| | **Reference** | **EviAnn** | **FINDER** | **MAKER2** | **BRAKER3** | **StringTie** |
| Genes | 27199 | 26928 | 33023 | 24520 | 28491 | 22293 |
| All transcripts | 35145 | 41161 | 35835 | 25962 | 31842 | 63442 |
| lncRNA transcripts | 0 | 201 | 25885 | 0 | 0 | |
| Unique proteins | 32609 | 35108 | 9145 | 25383 | 30846 | |
| Species | *D. melanogaster* | | | | | |
| | **Reference** | **EviAnn** | **FINDER** | **MAKER2** | **BRAKER3** | **StringTie** |
| Genes | 15986 | 12810 | 29124 | 11762 | 14057 | 15007 |
| All transcripts | 35145 | 26594 | 31564 | 13830 | 17239 | 49171 |
| Long non-coding RNA transcripts | 2258 | 315 | 23336 | 0 | 0 | |

| | | | | | | |
|---|---|---|---|---|---|---|
| Unique proteins | 21473 | 19188 | 6940 | 13160 | 16721 | |
| Species | *P. trichocarpa* | | | | | |
| | **Reference** | **EviAnn** | **FINDER** | **MAKER2** | **BRAKER3** | **StringTie** |
| Genes | 31697 | 30620 | 43211 | 39645 | 35088 | 31997 |
| All transcripts | 58718 | 56220 | 54318 | 43070 | 40737 | 103729 |
| lncRNA transcripts | 7078 | 542 | 23158 | 0 | 0 | |
| Unique proteins | 43029 | 45734 | 26288 | 41875 | 39630 | |
| Species | *D. rerio* | | | | | |
| | **Reference** | **EviAnn** | **FINDER** | **MAKER2** | **BRAKER3** | **StringTie** |
| Genes | 38655 | 27054 | 31164 | | 25788 | 46958 |
| All transcripts | 66521 | 42819 | 33947 | | 30705 | 126980 |
| lncRNA transcripts | 9962 | 650 | 21963 | | 0 | |
| Unique proteins | 50045 | 39926 | 11837 | | 29990 | |
| Species | *G. gallus* | | | | | |
| | **Reference** | **EviAnn** | **FINDER** | **MAKER2** | **BRAKER3** | **StringTie** |
| Genes | 24376 | 16291 | 45596 | 17995 | 24242 | 24089 |
| All transcripts | 83358 | 46726 | 51441 | 24232 | 30238 | 114588 |
| lncRNA transcripts | 14238 | 502 | 14238 | 0 | 0 | |
| Unique proteins | 46607 | 35906 | 13325 | 23042 | 29634 | |
| Species | *M. musculus* | | | | | |
| | **Reference** | **EviAnn** | **FINDER** | **MAKER2** | **BRAKER3** | **StringTie** |
| Genes | 25906 | 21260 | 28913 | | 20991 | 27068 |
| All transcripts | 58133 | 46521 | 32790 | | 26925 | 99421 |
| lncRNA transcripts | 8295 | 240 | 19317 | | 0 | |
| Unique proteins | 38688 | 39771 | 12508 | | 26476 | |

| Supplementary Table 9. Gffcompare codes for annotations of transcripts for EviAnn | | | | | | |
|---|---|---|---|---|---|---|
| Species | *A. thaliana* | | | | | |
| Code | **= (complete match)** | **j (at least one junction matches)** | **c (contains the reference)** | **m (retained introns)** | **k (contained in the reference)** | **u (no overlap with anything in the reference)** |
| Species | *A.thaliana* | | | | | |
| EviAnn transcripts | 24599 | 8946 | 1894 | 1919 | 1059 | 1248 |
| Species | *D. melanogaster* | | | | | |
| EviAnn transcripts | 15129 | 6684 | 1533 | 1208 | 1113 | 244 |
| Species | *P. trichocarpa* | | | | | |
| EviAnn transcripts | 30264 | 13871 | 3030 | 3641 | 1046 | 1750 |
| Species | *D.rerio* | | | | | |
| EviAnn transcripts | 21193 | 11095 | 6712 | 704 | 319 | 1690 |
| Species | *G. gallus* | | | | | |
| EviAnn transcripts | 21864 | 17122 | 3430 | 1273 | 1575 | 413 |
| Species | *M. musculus* | | | | | |
| EviAnn transcripts | 21818 | 16325 | 4913 | 832 | 472 | 1303 |

**Command lines used to run the pipelines and for evaluating the results.**

Substitute genome.fna and protein.faa with the appropriate file names for files containing genome and protein sequences.

EviAnn: eviann.sh -t 16 -g genome.fna -r rnaseq.txt -p protein.faa

BRAKER3: singularity exec braker3.sif braker.pl –genome=genome.fna --prot_seq=protein.faa --bam= RNA-Seq.bam --workingdir=work_dir --threads=16 --softmask

Maker2: ez_maker.sh -t 16 -g genome.fna -r rnaseq.txt -p protein.faa -d

FINDER: singularity exec --disable-cache -B /data:/data /ccb/sw/packages/Finder-finder_v1.1.0/finder.sif finder --genome genome.fna --protein protein.faa --metadatafile SRR.metadata --output_directory out_dir --organism_model VERT --cpu 16 --verbose 1 -no_cleanup --checkpoint 0 --genemark_path /ccb/sw//packages/GeneMark-ETP/bin//gmes/ --genemark_license ~/.gm_key

**Evaluation script (also available on github under eviann/ submodule as evaluate.sh):**

```
$ cat evaluate.sh

#!/bin/bash
#this code evaluates sensitivity and precision of genes and transcripts counting
either a transcript or a CDS match as a match
#!/bin/bash
#this code evaluates sensitivity and precision of genes and transcripts counting
either a transcript or a CDS match as a match
function usage {
  echo "Usage: evaluate_gffcompare.sh [options]"
  echo "Options:"
  echo " -r FILE    reference annotation in GFF format"
  echo " -q FILE    query annotation in GFF format"
  echo " -g FILE    genome sequence file in fasta format"
}

while [[ $# > 0 ]]
do
    key="$1"

    case $key in
        -r|--reference)
            REF="$2"
            shift
            ;;
        -q|--query)
            QRY="$2"
            shift
```

```
            ;;
        -g|--genome)
            GENOME="$2"
            shift
            ;;
        -h|--help|-u|--usage)
            usage
            exit 255
            ;;
        *)
            echo "Unknown option $1"
            usage
            exit 1        # unknown option
            ;;
    esac
    shift
done

if [ ! -s $REF ] || [ ! -s $QRY ] || [ ! -s $GENOME ];then
  echo "ERROR: one of the input files is empty of does not exist"
  exit 1
fi

REFL=`basename $REF`
QRYL=`basename $QRY`

#compute loci in the query
gffread -M --cluster-only -F --keep-genes $QRY > $QRYL.fix

#filter the refseq annotation keeping only protein coding and lnc_RNA genes
awk -F '\t' '{if($7 != "?") print }' $REF | \
  gffread -M -F -J -g $GENOME --ids <(perl -F'\t' -ane '{if(($F[2] eq "lnc_RNA" ||
$F[2] eq "mRNA" || $F[2] eq "transcript" || $F[2] eq
"primary_transcript")){unless($F[8] =~/pseudo=true|exception=dicistronic
gene|exception=trans-splicing|exception=RNA editing/){@f=split(";",$F[8]);print
substr($f[0],3),"\n"}}}' $REF ) > $REFL.gene_transcript_lncRNA.gff

#produce table of locus gene transcript
perl -F'\t' -ane '{if($F[2] eq "lnc_RNA" || $F[2] eq "mRNA" || $F[2] eq "transcript"
|| $F[2] eq "primary_transcript"){@f=split(/;/,$F[8]);print "$locus
",substr($f[1],7)," ",substr($f[0],3),"\n";}elsif($F[2] eq
"locus"){@f=split(/;/,$F[8]);$locus=substr($f[0],3);}}'
$REFL.gene_transcript_lncRNA.gff > $REFL.locus_gene_transcript

perl -F'\t' -ane '{if($F[2] eq "lnc_RNA" || $F[2] eq "mRNA" || $F[2] eq "transcript"
|| $F[2] eq "primary_transcript"){@f=split(/;/,$F[8]);print "$gene
",substr($f[0],3),"\n";}elsif($F[2] eq
"gene"){@f=split(/;/,$F[8]);$gene=substr($f[0],3);}}' $QRYL.fix >
$QRYL.gene_transcript

#produce CDS-only reference
```

```
gffread -C $REFL.gene_transcript_lncRNA.gff | \
  perl -F'\t' -ane '{unless($F[2] eq "exon"){if($F[2] eq "CDS"){$F[2]="exon"}print
join("\t",@F);}}' |\
  gffread -M > $REFL.gene_transcript_lncRNA.CDSasEXONS.gff

#produce CDS-only query
gffread -C $QRYL.fix|\
  perl -F'\t' -ane '{unless($F[2] eq "exon"){if($F[2] eq "CDS"){$F[2]="exon"}print
join("\t",@F);}}' |\
  gffread -M > $QRYL.fix.CDSasEXONS.gff

#compute which transcript match
cat <(trmap -c '=' $REFL.gene_transcript_lncRNA.gff $QRYL.fix) |\
  perl -ane
'{if($F[0]=~/^>/){$transc=substr($F[0],1)}else{$h{$F[5]}=$transc}}END{open(FILE,"'$RE
FL'.locus_gene_transcript");while($line=<FILE>){chomp($line); @f=split(/\s/,$line);
print $line," $h{$f[2]}\n" if(defined $h{$f[2]});}}' >
$REFL.matched_locus_gene_transcript

#compute which CDSs match
cat <(trmap -c '=' --strict-match $REFL.gene_transcript_lncRNA.CDSasEXONS.gff
$QRYL.fix.CDSasEXONS.gff) |\
  perl -ane
'{if($F[0]=~/^>/){$transc=substr($F[0],1)}else{$h{$F[5]}=$transc}}END{open(FILE,"'$RE
FL'.locus_gene_transcript");while($line=<FILE>){chomp($line); @f=split(/\s/,$line);
print $line," $h{$f[2]}\n" if(defined $h{$f[2]});}}' > $REFL.matched_locus_gene_CDS

#produce matched qry loci
cat $REFL.matched_locus_gene_transcript $REFL.matched_locus_gene_CDS |\
  perl -ane
'{$h{$F[3]}=1}END{open(FILE,"'$QRYL'.gene_transcript");while($line=<FILE>){chomp($lin
e);@f=split(/\s/,$line);print $line,"\n" if(defined $h{$f[1]});}}' >
$QRYL.matched_locus_transcriptORCDS

#compute statistcs
N_REF_TRANSCRIPTS=`awk '{print $3}' $REFL.locus_gene_transcript | wc -l`
N_REF_GENES=`awk '{print $2}' $REFL.locus_gene_transcript | sort|uniq |wc -l`
N_MATCHING_REF_TRANSCRIPTS=`awk '{print $3}' $REFL.matched_locus_gene_transcript
|sort |uniq |wc -l`
N_MATCHING_REF_CDS=`awk '{print $3}' $REFL.matched_locus_gene_CDS |sort |uniq |wc -l`
N_MATCHING_REF_GENES=`cat $REFL.matched_locus_gene_transcript
$REFL.matched_locus_gene_CDS | awk '{print $2}' |sort |uniq |wc -l`
N_REF_CDS=`awk -F '\t' '{if($3=="transcript" ||$3=="mRNA"|| $3=="lnc_RNA" ||
$3=="primary_transcript") print}' $REFL.gene_transcript_lncRNA.CDSasEXONS.gff |wc -l`

N_QRY_TRANSCRIPTS=`awk -F '\t' '{if($3=="transcript" ||$3=="mRNA"|| $3=="lnc_RNA")
print}' $QRYL.fix  |wc -l`
N_QRY_LOCI=`awk '{print $1}' $QRYL.gene_transcript |sort|uniq|wc -l`
N_MATCHING_QRY_LOCI=`awk '{print $1}' $QRYL.matched_locus_transcriptORCDS |sort| uniq
| wc -l`
```

```
N_MATCHING_QRY_TRANSCRIPTS=`awk '{print $4}' $REFL.matched_locus_gene_transcript
|sort |uniq |wc -l`
N_MATCHING_QRY_CDS=`awk '{print $4}' $REFL.matched_locus_gene_CDS |sort |uniq |wc -l`
N_QRY_CDS=`awk -F '\t' '{if($3=="transcript" ||$3=="mRNA"|| $3=="lnc_RNA") print}'
$QRYL.fix.CDSasEXONS.gff |wc -l`

#compute Sn Pr and F1 and print
awk 'BEGIN{print "Transcript/CDS and gene evaluations:";
  print "Reference Transcripts: '$N_REF_TRANSCRIPTS' Reference genes:
'$N_REF_GENES'";
  tsn=int('$N_MATCHING_REF_TRANSCRIPTS'/'$N_REF_TRANSCRIPTS'*1000+.5)/10;
  tpr=int('$N_MATCHING_QRY_TRANSCRIPTS'/'$N_QRY_TRANSCRIPTS'*1000+.5)/10;
  csn=int('$N_MATCHING_REF_CDS'/'$N_REF_CDS'*1000+.5)/10;
  cpr=int('$N_MATCHING_QRY_CDS'/'$N_QRY_CDS'*1000+.5)/10;
  gsn=int('$N_MATCHING_REF_GENES'/'$N_REF_GENES'*1000+.5)/10;
  gpr=int('$N_MATCHING_QRY_LOCI'/'$N_QRY_LOCI'*1000+.5)/10;
  print "Correct transcripts: '$N_MATCHING_QRY_TRANSCRIPTS' Matched reference
transcripts: '$N_MATCHING_REF_TRANSCRIPTS' Total query transcripts:
'$N_QRY_TRANSCRIPTS'\nCorrect CDS: '$N_MATCHING_QRY_CDS' Matched Reference CDS:
'$N_MATCHING_REF_CDS' Total unique query CDS: '$N_QRY_CDS' Total unique reference
CDS: '$N_REF_CDS'\nCorrect gene loci: '$N_MATCHING_QRY_LOCI' Total query gene loci:
'$N_QRY_LOCI'\nTranscript Sensitivity: "tsn" Precision: "tpr" F1:
"int(2*tsn*tpr/(tsn+tpr)*10+0.5)/10"\nCDS Sensitivity: "csn" Precision: "cpr" F1:
"int(2*csn*cpr/(csn+cpr+0.001)*10+0.5)/10"\nGene Sensitivity: "gsn" Precision: "gpr"
F1: "int(2*gsn*gpr/(gsn+gpr)*10+.5)/10}'
```
